# The *Sw-5b* NLR immune receptor induces earlier transcriptional changes in response to thrips-mediated inoculation of *Tomato spotted wilt orthotospovirus* compared to mechanical inoculation

**DOI:** 10.1101/2022.09.07.507022

**Authors:** Norma A. Ordaz, Ugrappa Nagalakshmi, Leonardo S. Boiteux, Hagop S. Atamian, Diane E. Ullman, Savithramma P. Dinesh-Kumar

## Abstract

The nucleotide-binding leucine-rich repeat (NLR) class of immune receptor, *Sw-5b* confers resistance to *Tomato spotted wilt orthotospovirus* (TSWV). Although *Sw-5b* is known to activate immunity upon recognition of the NSm of TSWV, we know very little about the downstream events that lead to resistance. Here, we investigated the early transcriptomic changes that occur in response to both mechanical and thrips-mediated inoculation of TSWV using near-isogenic resistant and susceptible tomato lines. Interestingly, the *Sw-5b* induces earlier transcriptional changes in response to thrips-mediated inoculation compared to mechanical inoculation of TSWV. A subset of the differentially expressed genes (DEGs) observed at 12 and 24 hours post thrips-mediated inoculation of TSWV was only present at 72 hours post mechanical inoculation. Although some DEGs were shared between thrips and mechanical inoculation at 72 hours postinfection, many DEGs were specific to either thrips-mediated or mechanical inoculation of TSWV. In response to thrips-mediated inoculation, an NLR immune receptor, cysteine-rich receptor-like kinase, G-type lectin S-receptor-like kinases, and transcription factors such as the ethylene response factor 1 and the calmodulin-binding protein 60 were induced. Whereas, in response to mechanical inoculation, fatty acid desaturase 2-9, cell death genes, DCL2b, RIPK/PBL14-like, and transcription factors such as ERF017 and WRKY75 were differentially expressed. Our findings reveal novel insights into *Sw-5b* responses specific to the method of TSWV inoculation. Given that TSWV is transmitted in nature primarily by the thrips, the DEGs we have identified provide a foundation for understanding the mechanistic roles of these genes in the *Sw-5b*-mediated resistance.

## INTRODUCTION

*Tomato spotted wilt orthotospovirus* (TSWV) is one of the most economically important plant viruses affecting tomato production worldwide (Oliver and Whitfield, 2016; Zhu et al., 2019). TSWV is the type member of the genus *Orthotospovirus*, family *Tospoviridae*, order *Bunyavirales*, and is characterized by a single-stranded negative-sense RNA genome composed of three RNA segments enclosed in a host-derived virion envelope with two embedded glycoproteins (Zhu et al., 2019). TSWV can be transmitted mechanically with infected sap (Rotenberg et al., 2015); however, in nature, the western flower thrips (WFT), *Frankliniella occidentalis* (Pergande) primarily transmits orthotospoviruses. Transmission by the insect is in a circulative-propagative manner, and the orthotospoviruses ultimately invade and replicate in the foregut, midgut, tubular, and principal salivary glands of the vector, from which they are injected into plants during feeding (Ullman et al., 1993a; Ullman et al., 1993b; Ullman et al., 1995; Whitfield et al., 2005; Montero-Astua et al., 2016).

Several TSWV resistance genes (*Sw-1a, Sw-1b, Sw-2, Sw-3, Sw-4, Sw-5, Sw-6*, and *Sw-7*) have been described in tomatoes (Brommonschenkel et al., 2000; Dianese et al., 2011) With regards to orthotospovirus resistance, the most effective among them is *Sw-5*, a single dominant resistance gene locus that was introgressed from *Solanum peruvianum* to a commercial tomato line ‘Stevens’ (Stevens, 1964). The gene was mapped to the end of chromosome nine (Brommonschenkel et al., 2000; de Oliveira et al., 2018). Within the *Sw-5* gene cluster, *Sw-5a* has been shown to be nonfunctional against TSWV (Spassova et al., 2001; Hallwass et al., 2014; De Oliveira et al., 2016), but was recently shown to provide resistance against the geminivirus, *Tomato leaf curl New Delhi virus*, by recognizing an AC4 protein (Sharma et al., 2021). *Sw-5b* has been shown to mediate resistance against TSWV, and several other tomato-infecting orthotospoviruses (Boiteux and Giordano, 1993; Spassova et al., 2001; De Oliveira et al., 2016; de Oliveira et al., 2018). The *Sw-5b* gene encodes a coiled-coil nucleotide-binding leucine-rich repeat (CC-NLR) class of immune receptors (Brommonschenkel et al., 2000; Zhu et al., 2019). In addition, *Sw-5b* contains an extended N-terminal Solanaceae domain (SD) that is only present in some CC-NLRs from solanaceous species (Chen et al., 2016).

The *Sw-5b* NLR recognizes the cell-to-cell movement protein (MP), NSm, of TSWV and induces a localized cell death response called the hypersensitive response (HR) at the site of infection to limit virus spread (Hallwass et al., 2014; Peiro et al., 2014). Within NSm, Sw-5b specifically recognizes a conserved 21 amino acid motif (NSm^21^), and confers resistance against most of the American-type orthotospoviruses (Zhu et al., 2017). Interestingly, *Sw-5b* adopts a unique two-step recognition mechanism to induce a robust immune response against TSWV (Li et al., 2019). In addition to the LRR domain recognizing NSm and NSm^21^, the N-terminal SD also directly interacts with NSm, and this interaction is critical for recognition. In the absence of TSWV NSm, the CC domain keeps the NB-LRR region of Sw-5b in an autoinhibited state to avoid induction of an autoimmune response (Chen et al., 2016). The recognition of NSm by SD releases the autoinhibition state leading to activation of the NB-LRR region, and induction of HR cell death. The SD also facilitates recognition of low levels of NSm by the NB-LRR domain of Sw-5b to activate immunity. Therefore, Sw-5b uses two domains to perceive the viral effector NSm, and to activate a robust defense against TSWV (Li et al., 2019).

Although we have some insights on how *Sw-5b* NLR recognizes the viral effector NSm, we know very little about the mechanisms involved immediately downstream of effector recognition, especially the transcriptional changes that activate immune responses leading to cell death and containment of virus to the infection site. In addition, there is no report of gene expression changes that occur in *Sw-5b* plants in response to TSWV infection with thrips, even though this is the primary manner of spread in the field. Therefore, we investigated the early transcriptional changes that occur during *Sw-5b* NLR recognition of TSWV in tomato when infected with TSWV via thrips and compared it with TSWV infection through mechanical inoculation. We found that the *Sw-5b* NLR induces earlier transcriptomic changes in response to thrips-mediated inoculation of TSWV compared to mechanical inoculation, and that many differentially expressed genes (DEGs) were specific to either thrips-mediated or mechanical inoculation of TSWV. Our findings provide new insight into genes that could play a role in *Sw-5b*-mediated resistance to TSWV and highlight the importance of understanding the *Sw-5b*-mediated signaling pathway when activated by thrips inoculation.

## RESULTS AND DISCUSSION

### Optimization of TSWV inoculation and tissue collection for transcriptomic analysis

We used the near-isogenic tomato lines, ‘Santa Clara’ (*Sw-5b^-/-^*, the TSWV-susceptible homozygous line) and ‘CNPH-LAM 147’ with *Sw-5b (Sw-5b^+/+^*, the TSWV-resistant homozygous line) (Dianese et al., 2010; Hallwass et al., 2014) for investigating the transcriptomic changes that occur during *Sw-5b*-mediated resistance to TSWV infection. Since robust infection with TSWV is important to induce *Sw-5b* responses, we first optimized the developmental stage (DS) of plants that would best support virus infection using susceptible ‘Celebrity’ tomato plants, and the MR-01 strain of TSWV (TSWV^MR-01^). We rub-inoculated a known amount of sap prepared from TSWV^MR-01^ infected *Datura stramonium* tissue (hereafter referred to as mechanical inoculation) onto leaves of tomato plants at the 2, 3, 4, 5, 6, and 7 leaf stage (DS2 to DS7) (Fig. 1A). Nearly 100% of DS2 to DS4 stage ‘Celebrity’ tomato plants were systemically infected as measured by enzyme linked-immunosorbent assay (ELISA) and developed symptoms by 12 days postinoculation (Fig. 1B). The percentage of plants showing symptoms decreased when inoculation was done at later developmental stages (Fig. 1B). Only about 50% and 10% of the plants at DS6 and DS7 respectively, were symptomatic (Fig. 1B). Regression analysis showed a positive and statistically significant (p=0.0004) correlation between developmental stage and symptom expression (Fig. 1C). Based on these results we used DS2 stage ‘Santa Clara’ and ‘CNPH-LAM 147’ plants for mechanical inoculation with TSWV^MR-01^ using our optimized conditions (see materials and methods section for details) to prepare tissue for RNA-seq. During those experiments we collected the two inoculated leaves from each plant at 0, 12, 24 and 72-hours post-inoculation from three biological replicates (Fig. 1D).

**Figure 1.**
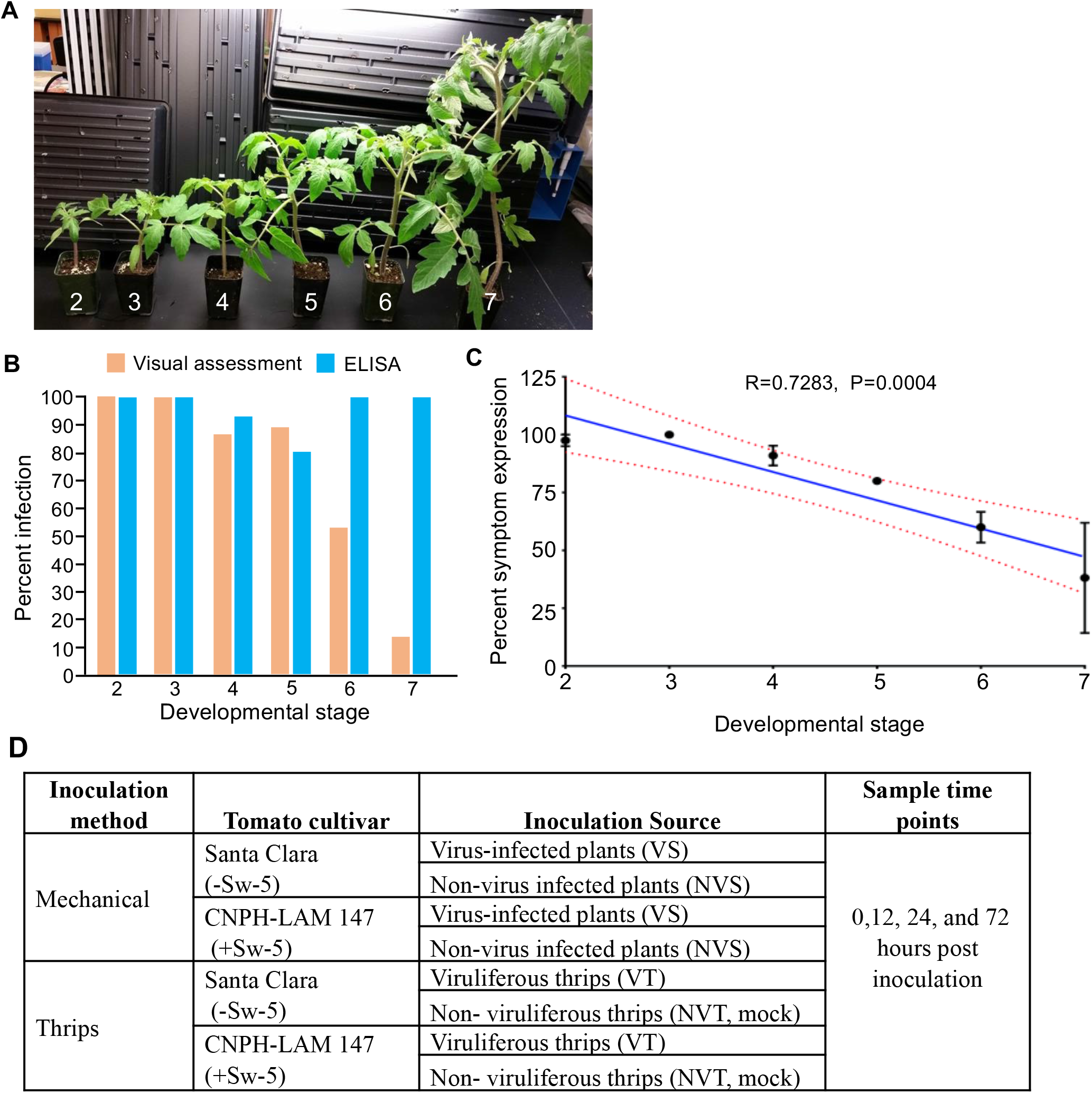
Virus inoculation and tissue collection scheme used for RNA-seq analysis. (A)Plants at six developmental stages (DS) used for mechanical inoculation of TSWV to determine the optimal DS for tissue collection for RNA-seq experiments. (B) Comparison of mean percent infection based on visual assessment to ELISA of susceptible Celebrity tomato 12 days post mechanical inoculation (dpi) of TSWV at six developmental stages. (C) Correlation between plant developmental stage and symptom expression at 12 dpi (regression analysis, mean percent of plants with symptoms). Data in B and C are from two experimental replicates (rep 1, n=20-21 plants/DS and rep 2, n = 5 to 15 plants/DS). (D) Schematics of treatments used for RNA-seq experiments. Isogenic susceptible tomato line Santa Clara and resistant tomato line CNPH-LAM 147 were inoculated with TSWV by mechanical or through thrips and tissue samples were collected at 0, 12, 24 and 72 hours post inoculation (HPI).

Thrips transmission involves three phases: acquisition, latent period, and inoculation. Importantly, acquisition only occurs during the larval stages, with inoculation occurring primarily during the adult stage. To optimize the transmission process, we standardized the age of larvae used for acquisition of TSWV, determining that allowing TSWV acquisition by first instar larvae less than six-hours post eclosion increased the probability of TSWV inoculation by adults to 80% to 100%. The optimum number of thrips to attain this level of inoculation was ten thrips (5 females and 5 males). During experiments to prepare materials for RNA-seq, thrips were placed on leaf 1 and leaf 2 of each DS2 stage tomato plant (‘Santa Clara’ and ‘CNPH-LAM 147’) that had been placed inside a closed cage (Supplementary Fig. S1; see materials and methods section for details). Thrips remained on the plants until leaf samples were collected at 0, 12, 24 and 72-hours post-thrips placement from three biological replicates (Supplementary Fig. S1). Total RNA was extracted, and RNA-seq libraries were prepared. Illumina sequencing was performed as described in the materials and methods section.

### Overview of transcriptomic changes observed in *Sw-5b* resistant plants in response to TSWV infection

A total of 1.1 billion 50 bp reads were generated using Illumina sequencing. The number of reads per sample ranged from 5 to 30 million. After quality control and filtering low quality reads, 84% of the reads were mapped to the reference tomato cDNA sequences (version ITAG4.1; https://solgenomics.net/). The mapped reads ranged from 4.5 to 25 million reads per sample. The general linear model of the edgeR package in R was used to identify the differentially expressed genes (DEGs) in the ‘CNPH-LAM 147’ resistant near-isogenic line compared to the susceptible ‘Santa Clara’ for each inoculation method and time point. Mock inoculation with sap prepared from non-infected plants or non-infected thrips were included as controls. These controls were used to observe base gene expression and identify whether any DEGs from TSWV inoculated treatments were being expressed in the mock inoculations with sap prepared from non-infected plants or inoculation with non-infected thrips. These comparisons allowed removal of any DEGs activated by mechanical wounding and non-infected thrips feeding damage from our analysis. Among these experimental controls, we found no differences between resistant and susceptible lines using the general linear model. Consequently, the differences in gene expression we detected in the resistant line are specifically due to the interaction between TSWV and *Sw-5b*. Genes with more than two-fold expression difference (false discovery rate < 0.05) were considered significantly different. We identified a total of 80 and 95 DEGs in the resistant line across three different time points post-thrips and mechanical inoculation, respectively (Fig. 2A and Supplementary Table S1). Of the 175 DEGs identified, 136 were upregulated and 35 were downregulated (Fig. 2A and Supplementary Table S1).

**Figure 2.**
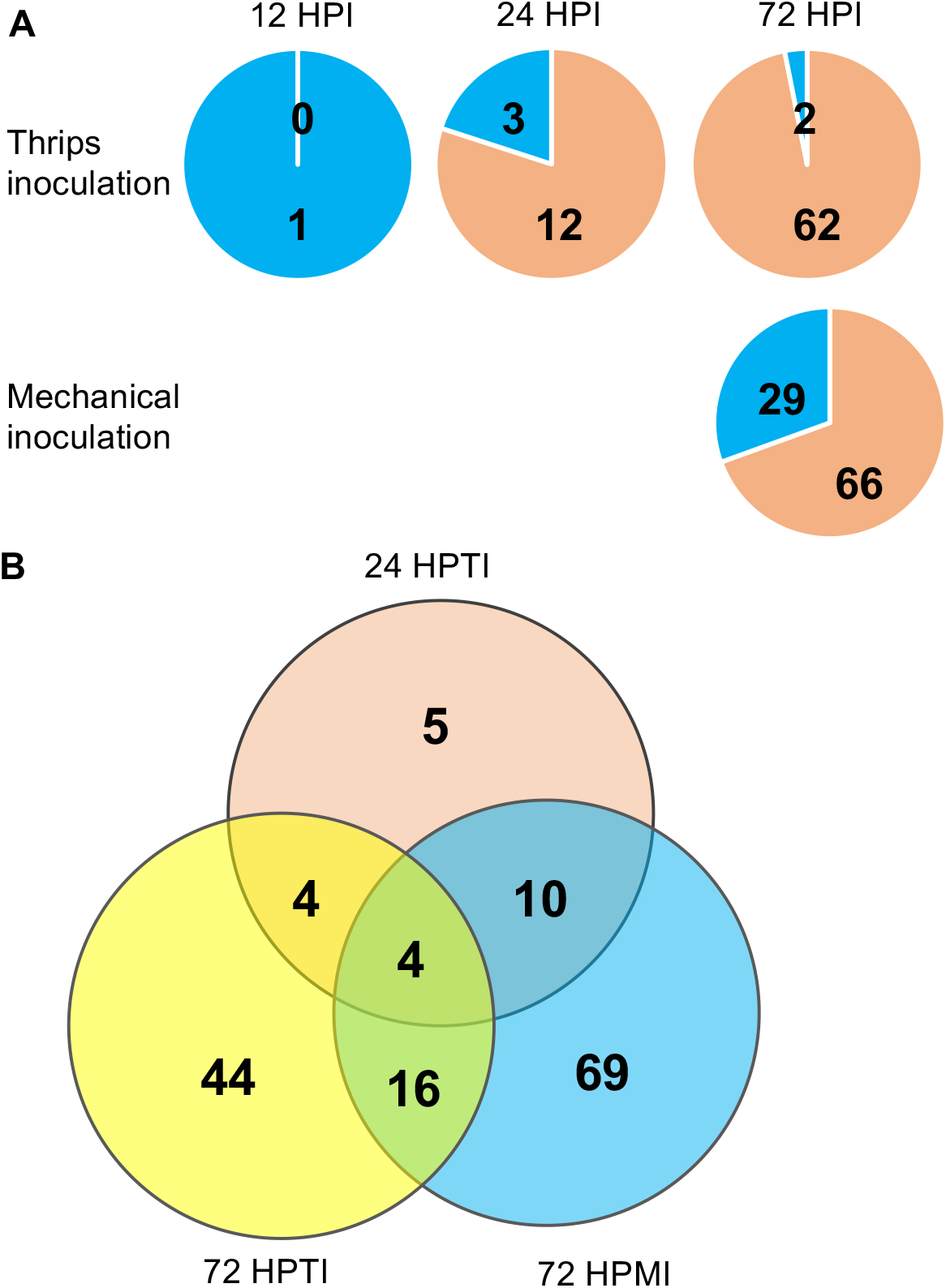
Venn diagrams showing the number of differentially expressed genes (DEGs). (A)Number of DEGs detected at 12, 24 and 72 hours post thrips inoculation (top panels) and mechanical inoculation (bottom panel) of TSWV in *Sw-5b* resistant tomato line compared to susceptible line. Orange, upregulated DEGs; blue, down-regulated DEGs. (B) Number of unique and common DEGs detected among different time points and different inoculation methods. HPTI, hours post thrips-mediated inoculation of TSWV; HPMI, hours post mechanical inoculation of TSWV; HPI, hours post-inoculation.

### The *Sw-5b* NLR induces early transcriptomic changes to thrips-mediated inoculation of TSWV

We observed clear differences in timing of transcriptomic changes between the inoculation methods. In thrips inoculated plants, DEGs were detected as early as 12 and 24 hours post-thrips inoculation (HPTI) (Fig. 2A and Supplementary Table S1). Whereas, in mechanically inoculated plants, DEGs were not detected until 72 hours post-mechanical inoculation (HPMI). At 12 HPTI we observed a single downregulated gene. At 24 HPTI, 12 genes were upregulated and three genes were downregulated. Among the 64 DEGs identified at 72 HPTI, 62 were upregulated and 2 were downregulated. In 72 HPMI plants, 66 genes were upregulated and 29 genes were downregulated. Only 4 DEGs were shared between two inoculation methods and different time points (Fig. 2B and Table 1). Ten genes that were differentially expressed at 24 HPTI were only observed at the later time point for mechanical inoculation (72 HPMI) (Fig. 2B and Table 1). A significant number of DEGs at 72 hours post infection were specifically expressed either in the thrips inoculated or mechanically inoculated plants (Fig. 2B and Supplementary Table S1). These results indicated that thrips inoculation induced earlier *Sw-5b* NLR-mediated transcriptome changes, and that many of the DEGs induced were unique to inoculation method.

**Table 1.** Differentially expressed genes in *Sw-5b* resistant plant at 24 hour post thrips inoculation of TSWV that are shared with 72 hours post-thrips and mechanical inoculation.

| Gene ID | Thrips Inoculation |  | Mechanical Inoculation | Annotation |
| --- | --- | --- | --- | --- |
|  | 24 hpi | 72 hpi | 72 hpi |  |
|  | logFC | logFC | logFC |  |
| Solyc03g025670 | 4.55 | 3.70 | 5.19 | Pathogenesis-related protein 1c |
| Solyc01g097240 | 4.13 | 4.48 | 6.40 | Pathogenesis-related protein 4 |
| Solyc09g006005 | 3.58 | 3.48 | 5.74 | Pathogenesis-related protein 1 |
| Solyc10g079860 | 2.81 | 2.23 | 4.68 | Beta(1,3) glucanase |
| Solyc01g101180 | 6.33 |  | 5.00 | Terpene synthase |
| Solyc02g070110 | 4.12 |  | 3.88 | FAD-binding Berberine family protein |
| Solyc10g011925 | 2.63 |  | 3.14 | Phenylalanine ammonia-lyase |
| Solyc03g115060 | 2.48 |  | 2.54 | SRP40-like protein |
| Solyc10g076510 | 2.25 |  | 2.22 | Pyruvate decarboxylase |
| Solyc03g123540 | -3.01 |  | -3.09 | Class III heat shock protein - HSP17.4 |
| Solyc05g051350 | 8.59 |  |  | Rhamnogalacturonate lyase |
| Solyc09g091670 | 3.95 |  |  | Pleiotropic drug resistance protein |
| Solyc11g010210 | 2.52 |  |  | WEB family protein |
| Solyc03g013340 | -1.23 |  |  | NOD26-like intrinsic protein 2.1 |
| Solyc03g007890 | -1.50 |  |  | Class II heat shock protein -HSP17.6 |

A single downregulated DEG (Solyc09g082340) (−8.18 Log fold) at 12 HPTI (Supplementary Table S1) is predicted to encode a vicilin-like protein. Vicilin was not differentially expressed at later time points in thrips inoculated treatments nor in mechanically inoculated treatments, suggesting the importance of this gene in early *Sw-5b*-mediated resistance to thrips inoculation of TSWV^MR-01^. Vicilins are plant-specific proteins and structurally belong to the cupin superfamily of proteins (Dunwell et al., 2001; Chen et al., 2013). Structure-guided biochemical analysis of vicilin from pepper and tomato indicate that vicilin may function as superoxide dismutase (SOD) to regulate oxidative stress (Shikhi et al., 2018; Shikhi et al., 2020). Furthermore, the C-terminal cupid fold of pepper and tomato vicilin in the crystal structure is bound by the defense hormone salicylic acid (SA), suggesting that SA could modulate the SOD activity of vicilin (Shikhi et al., 2018; Shikhi et al., 2020). Given that reactive oxygen species (ROS) play an important early defense role (Castro et al., 2021), *Sw-5b*-mediated recognition of TSWV^MR-01^ could downregulate vicilin to suppress SOD activity. In *Arabidopsis, PAP85* encodes a vicilin-like protein that is upregulated early during *Tobacco mosaic virus* (TMV) infection, and virus accumulation was reduced in *AtPAP85* RNAi plants (Chen et al., 2013). Since TMV replication was reduced in protoplasts prepared from RNAi plants, it was proposed that *AtPAP85* might be involved in virus replication by facilitating endoplasmic reticulum (ER) membrane transition that is important for TMV replication. Interestingly, the NSm movement protein of TSWV that is recognized by *Sw-5b* to activate defense has been shown to physically interact with the ER membrane, and disruption of this interaction inhibits the cell-to-cell movement of NSm (Feng et al., 2016). Furthermore, in a *rdh3* mutant in which the ER network is altered, intracellular trafficking of NSm and systemic movement of TSWV were significantly affected (Feng et al., 2016). Therefore, we hypothesize that TSWV may use the vicilin-like protein for membrane interaction or reorganization to support virus replication and movement. Hence, *Sw-5b* as a defense strategy could downregulate vicilin early during the defense response to limit TSWV to the infection site. It will be interesting to test in the future, if knockdown or knockout of vicilin enhances resistance to TSWV.

Among the 15 DEGs detected at 24 HPTI in the *Sw-5b* resistant line, five were uniquely detected in response to thrips inoculation at this time point and were not detected at 72 HPTI, nor at any mechanical inoculation time points (Fig. 2B, Table 1 and Supplementary Table S1). These five unique DEGs included rhamnogalacturonate lyase (Solyc05g051350), pleiotropic drug resistance protein (Solyc09g091670), WEB family protein (Solyc11g010210), NOD26-like intrinsic protein 2.1 (Solyc03g013340), and a class II heat shock protein HSP17.6 (Solyc03g007890) (Supplementary Table S1). Pleiotropic drug resistance protein and the NOD26-like intrinsic protein are known to function as transporters facilitating the movement of diverse molecules across membranes in response to stimuli and stresses (Wallace et al., 2006; Nuruzzaman et al., 2014; Xie et al., 2021). In tobacco, pleiotropic drug resistance proteins have been shown to be regulated by defense hormones jasmonic acid (JA) and salicylic acid (SA) suggesting a regulatory role for these proteins in plant defense (Xie et al., 2021). In summary, these DEGs are of special interest for further characterization because they were specifically observed only in the thrips inoculated treatments.

### Specific transcriptomic responses important to plant defense are shared among thrips and mechanical inoculations

Some of the DEGs detected at 24 HPTI were also detected in the 72 HPTI and 72 HPMI (Fig. 2B, Table 1, and Supplementary Table S1). Among the 15 DEGs identified at 24 HPTI, four of them were detected in 72 HPTI and 72 HPMI treatments (Table 1). These DEGs were upregulated and included four pathogenesis-related (PR) proteins (Table 1), suggesting an important role for these well-known defense responsive genes in *Sw-5b*-mediated resistance. The upregulation of these DEGs indicates that as early as 24 HPTI, the *Sw-5b* NLR had initiated the defense response. That these same genes are also induced at 72 HPMI (Table 1) supports their importance in the *Sw-5b*-mediated resistance response independent of the inoculation method.

Out of 15 DEGs detected at 24 HPTI, eleven DEGs were not detected at 72 HPTI (Table 1). Among these eleven DEGs, six DEGs were detected at 72 HPMI indicating that these DEGs are responding to TSWV either delivered through thrips or through mechanical inoculation (Table 1). The four upregulated DEGs that encode enzymes are known to play a role in defense and these include terpene synthase, pyruvate decarboxylase, phenylalanine ammonia lyase (PAL), and FAD-binding berberine family protein (Table 1). Overexpression of terpene synthase in maize has been shown to enhance resistance to a fungal pathogen (Chen et al., 2018). The upregulation of pyruvate decarboxylase is known to induce PR proteins (Tadege et al., 1998; Rojas et al., 2014). In lettuce and sunflower FAD-binding berberine family proteins and BBE-like enzymes are upregulated after infection (Daniel et al., 2017; Benedetti et al., 2018). Although isochorismate synthase 1 (ICS1) is a major player in pathogen-induced SA biosynthesis, PAL has also been implicated in SA biosynthesis (Ding and Ding, 2020). Knockout of all PAL genes in Arabidopsis resulted in 50% decrease in pathogen-induced SA accumulation and were more susceptible to *Pseudomonas syringae* infection (Huang et al., 2010). In addition, both ICS and PAL pathways contribute to SA biosynthesis in soybean (Shine et al., 2016). PAL is also required for the biosynthesis of many other secondary metabolites including lignin (Pascual et al., 2016).

Sixteen DEGs were shared with both thrips and mechanical inoculation methods at 72 hours (Table 2 and Supplementary Table S1). Many of the shared DEGs are indicators of activation of plant defense. Eight of these DEGs encode PR proteins (Table 2). We observed upregulation of two DEGs that encode 1-aminocyclopropane-1-carboxylic acid synthase 2 (ACC synthase 2; Solyc01g095080) and ACC oxidase 1 (ACO1; olyc07g049530) enzymes that participate in ethylene production (Table 2) (Houben and Van de Poel, 2019). We also observed upregulation of lipoxygenase (LOX; Solyc08g029000), which is usually associated with responses to wounding from mechanical damage or thrips feeding damage resulting in the synthesis of JA (Yan et al., 2013). However, our experimental controls included mock inoculations with sap prepared from non-infected plants, and inoculation with non-infected thrips, which allowed us to remove responses due to mechanical wounding and thrips feeding damage from our analysis. Hence, upregulation of the lipoxygenase observed in our data is specific to the *Sw-5b*-mediated resistance to TSWV infection. It is possible that induced LOX during *Sw-5b*-mediated defense could increase JA synthesis which is known to impart resistance against thrips. In pepper, LOX2 is involved in JA biosynthesis and induces defense against western flower thrips (Sarde et al., 2019). We did not measure resistance to thrips in our experiments; however, this finding suggests that *Sw-5b*-mediated resistance results in outcomes that could negatively impact thrips biology, as well as TSWV infection. We also observed an upregulation of arogenate dehydrogenase (Solyc09g011870) and polyphenol oxidase (Solyc02g078650), both known to play an integral part in coordinating ROS leading to hypersensitive cell death responses.

**Table 2.** Differentially expressed genes in *Sw-5b* resistant plants that are shared between thrips and mechanical inoculation at 72 hours post TSWV infection.

| Gene ID | Thrips<br>Inoculation | Mechanical<br>Inoculation | Annotation |
| --- | --- | --- | --- |
|  | logFC | logFC |  |
| Solyc05g054380 | 9.12 | 7.35 | Pathogenesis related protein STH-2 |
| Solyc02g078650 | 8.69 | 6.66 | Polyphenol oxidase |
| Solyc01g008620 | 8.46 | 6.86 | Glucan endo-1,3-beta glucosidase A precursor |
| Solyc09g007010 | 8.26 | 6.48 | Pathogenesis-related protein PR1a |
| Solyc04g064880 | 7.46 | 6.88 | Pathogen-related protein-like |
| Solyc01g095080 | 6.74 | 5.67 | 1-aminocyclopropane-1-carboxylic acid synthase 2 |
| Solyc01g106620 | 6.02 | 6.26 | PR1 |
| Solyc01g080570 | 5.78 | 5.32 | Inosine-uridine preferring nucleoside hydrolase family |
| Solyc08g029000 | 5.77 | 7.24 | Lipoxygenase |
| Solyc09g011870 | 5.33 | 4.49 | Arogenate dehydrogenase 2 |
| Solyc01g059965 | 5.32 | 5.76 | Glucan endo-1,3-beta-glucosidase B |
| Solyc04g072280 | 5.01 | 5.88 | Laccase |
| Solyc08g080650 | 4.78 | 5.06 | Pathogenesis related protein P23 |
| Solyc07g049530 | 4.56 | 3.61 | 1-aminocyclopropane-1-carboxylate oxidase 1 |
| Solyc02g082920 | 3.86 | 4.60 | Endochitinase (PR3) |
| Solyc10g055810 | 3.16 | 4.92 | Chitinase (PR3) |

### Genes differentially expressed in *Sw-5b* specifically in response to 72 hours post-thrips inoculation of TSWV

Of the 64 DEGs, 44 were detected specifically in response to thrips inoculation at 72 hours (Fig. 2A and Supplementary Table S1). Some of the interesting DEGs that are specifically regulated at 72 HPTI are shown in Table 3. We detected upregulation of a CC-NLR, a cysteine-rich receptorlike kinase (CRK) and three G-type lectin S-receptor-like kinases (G-LecRLK) (Table 3). Given the emerging evidence that the helper or other NLRs function downstream of sensor NLRs to activate immunity (van Wersch et al., 2020), it will be interesting to test in the future the precise role of the upregulated CC-NLR (Solyc04g007490) that we detected in response to thrips inoculation of *Sw-5b* plants. CRKs have been shown to play an important role in cell death and disease resistance (Bourdais et al., 2015; Yadeta et al., 2017; Saintenac et al., 2021). LecRLKs are important regulators of defense responses against various pathogens and pests (Sun et al., 2020). Overexpression of G-LecRLK has been shown to increase plant defense to biotrophic pathogens (Zhao et al., 2018). G-LecRLKs in *Nicotiana attenuata* have been demonstrated to be involved in recognition of insect feeding, and required to induce a full defense response against *Manduca sexta* (Gilardoni et al., 2011). Therefore, it is possible that *Sw-5b* regulated genes could play a role in defense against thrips.

**Table 3.** Some interesting differentially expressed genes in *Sw-5b* resistant plant specifically in response to 72 hours post-thrips inoculation of TSWV.

| Gene ID | logFC | Annotation |
| --- | --- | --- |
| <u>NLR and receptor-like kinases</u> |  |  |
| Solyc04g007490 | 1.44 | CC-NLR disease resistance protein |
| Solyc12g005720 | 4.80 | Cysteine-rich receptor-like protein kinase |
| Solyc04g077340 | 7.08 | G-type lectin S-receptor-like kinase |
| Solyc05g008310 | 2.20 | G-type lectin S-receptor like protein kinase |
| Solyc07g063770 | 1.83 | G-type lectin S-receptor-like kinase |
| <u>Transcription factors</u> |  |  |
| Solyc04g016000 | 5.56 | Heat shock transcription factor protein 8 |
| Solyc07g053140 | 4.84 | Zinc finger protein/CONSTANS-like protein |
| Solyc06g069760 | 3.48 | Dof zinc finger protein |
| Solyc05g051200 | 2.91 | Ethylene-responsive factor 1 |
| Solyc03g113960 | 3.26 | CBP60-like |
| <u>Proteases and protease inhibitor</u> |  |  |
| Solyc08g079900 | 7.11 | Subtilisin-like protease |
| Solyc11g022590 | 1.91 | Trypsin inhibitor-like protein precursor |
| Solyc01g079980 | -1.48 | Aspartyl protease family protein |
| <u>Other interesting genes</u> |  |  |
| Solyc01g105450 | 4.45 | ABC transporter G family |
| Solyc05g053600 | 4.15 | Pleiotropic drug resistance protein |
| Solyc01g099010 | 3.88 | GDSL esterase/lipase |
| Solyc08g078870 | 3.51 | Lipid-transfer protein |
| Solyc05g008220 | 2.62 | PADRE gene family |
| Solyc12g008960 | 2.38 | IQM1-like protein |
| Solyc02g081980 | 2.27 | Apyrase |
| Solyc03g112700 | 1.80 | Herbivore elicitor-regulated |
| Solyc07g065660 | 1.69 | Cellulose synthase-like protein E1 |
| Solyc07g054600 | 1.20 | F-box protein |

We detected upregulation of five transcription factors at 72 HPTI in *Sw-5b* plants. Among these ethylene response factors (ERFs) and members of calmodulin-binding protein 60 (CBP60) are important players in defense (Huang et al., 2016; Zheng et al., 2022). ERFs are known to bind GCC box and promote expression of some PR genes. The ERF1 that we detected in this study when overexpressed in tobacco has been shown to induce PR genes and ethylene responses (Huang et al., 2004). In Arabidopsis, CBP60a and CBP60b function as positive regulators of immunity and overexpression of CBP60b induces constitutive defense responses (Huang et al., 2021). It will be interesting to further characterize the precise roles of ERF1 and CBP60 in *Sw-5b*-mediated resistance.

Other classes of genes that were differentially regulated during thrips-mediated inoculation belong to the proteases and protease inhibitor family (Table 3). We detected significant upregulation of a gene encoding subtilisin-like protease (Solyc08g079900) (Table 3). Subtilisin proteases have an established role in plant immunity against different pathogens (Figueiredo et al., 2018). The gene Solyc11g022590 that is upregulated is predicted to encode trypsin inhibitor-like protein and shares significant homology with Arabidopsis gene At1g73260 that regulates cell death triggered in response to infection of *Pseudomonas syringae* expressing AvrB effector (Li et al., 2008). Interestingly, Solyc01g079980 that encodes aspartyl protease was downregulated in response to thrips-mediated inoculation indicating that it could play a negative regulatory role during *Sw-5b*-mediated resistance to TSWV.

We also observed upregulation of other interesting genes at 72 hours post-thrips mediated inoculation of TSWV (Table 3). The Arabidopsis homologs of Solyc01g105450 that encodes ABC transporter family protein and Solyc05g053600 that encodes pleiotropic drug resistance protein have been shown to transport cuticular lipids (McFarlane et al., 2010) and abscisic acid (ABA) (Kang et al., 2010), respectively. The lipid transfer protein (Solyc08g078870) homolog DRN1 (DISEASE RELATED NONSPECIFIC LIPID TRANSFER PROTEIN 1) in Arabidopsis is required for resistance against various phytopathogens (Dhar et al., 2020). The calmodulin binding protein (Solyc12g008960) homolog in Arabidopsis is known to bind catalase 2 and regulates JA biosynthesis (Lv et al., 2019).

### Genes differentially expressed in *Sw-5b* specifically in response to 72 hours post-mechanical inoculation of TSWV

At 72 HPMI, 69 of the 95 DEGs were specific to mechanical inoculation at 72 hours (Fig. 2 and Supplementary Table S1). Interestingly the number of genes that were downregulated (28 genes) was greater in 72 hours post-mechanical inoculation treatment than in any other treatment (Supplementary Table S1). Among five DEGs belonging to the transcription factor family, three were upregulated and two were down regulated (Table 4). Compared to ERF1 that is upregulated in thrips-inoculated treatment at 72 hours (Table 3), we detected upregulation of a different ERF family member, ERF017 in response to mechanical inoculation of TSWV in *Sw-5b* plants. The WRKY75 that we detected during *Sw-5b* mediated resistance is regulated at the epigenetic level in response to biotic and abiotic stresses in *Solanaceae* plants (Lopez-Galiano et al., 2018).

**Table 4.**
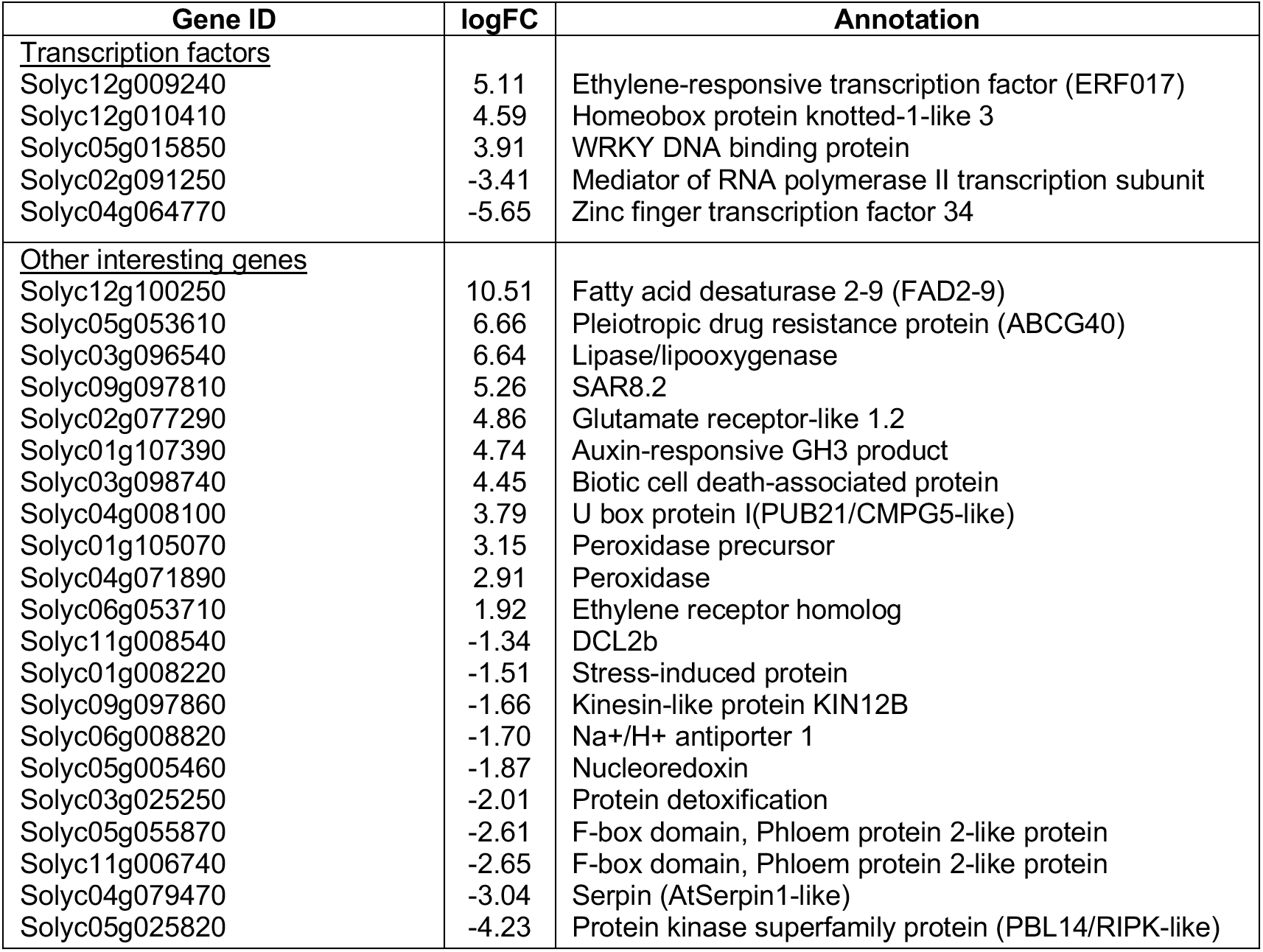
Some interesting differentially expressed genes in *Sw-5b* resistant plant specifically in response to 72 hours post-mechanical inoculation of TSWV.

| Gene ID | logFC | Annotation |
| --- | --- | --- |
| **Transcription factors** |  |  |
| Solyc12g009240 | 5.11 | Ethylene-responsive transcription factor (ERF017) |
| Solyc12g010410 | 4.59 | Homeobox protein knotted-1-like 3 |
| Solyc05g015850 | 3.91 | WRKY DNA binding protein |
| Solyc02g091250 | -3.41 | Mediator of RNA polymerase II transcription subunit |
| Solyc04g064770 | -5.65 | Zinc finger transcription factor 34 |
| **Other interesting genes** |  |  |
| Solyc12g100250 | 10.51 | Fatty acid desaturase 2-9 (FAD2-9) |
| Solyc05g053610 | 6.66 | Pleiotropic drug resistance protein (ABCG40) |
| Solyc03g096540 | 6.64 | Lipase/lipoxygenase |
| Solyc09g097810 | 5.26 | SAR8.2 |
| Solyc02g077290 | 4.86 | Glutamate receptor-like 1.2 |
| Solyc01g107390 | 4.74 | Auxin-responsive GH3 product |
| Solyc03g098740 | 4.45 | Biotic cell death-associated protein |
| Solyc04g008100 | 3.79 | U box protein I(PUB21/CMPG5-like) |
| Solyc01g105070 | 3.15 | Peroxidase precursor |
| Solyc04g071890 | 2.91 | Peroxidase |
| Solyc06g053710 | 1.92 | Ethylene receptor homolog |
| Solyc11g008540 | -1.34 | DCL2b |
| Solyc01g008220 | -1.51 | Stress-induced protein |
| Solyc09g097860 | -1.66 | Kinesin-like protein KIN12B |
| Solyc06g008820 | -1.70 | Na<sup>+</sup>/H<sup>+</sup> antiporter 1 |
| Solyc05g005460 | -1.87 | Nucleoredoxin |
| Solyc03g025250 | -2.01 | Protein detoxification |
| Solyc05g055870 | -2.61 | F-box domain, Phloem protein 2-like protein |
| Solyc11g006740 | -2.65 | F-box domain, Phloem protein 2-like protein |
| Solyc04g079470 | -3.04 | Serpin (AtSerpin1-like) |
| Solyc05g025820 | -4.23 | Protein kinase superfamily protein (PBL14/RIPK-like) |

Some of the other interesting genes regulated at 72 hours post-mechanical inoculation are shown in Table 4. The fatty acid desaturase 2-9 (FAD2-9) was significantly induced (10.51 Log fold). FADs are known to regulate ROS signaling and fluidity of cell membranes during defense (Xiao et al., 2022). Some genes involved in cell death such as the gene encoding biotic cell death associated protein (Solyc03g098740) was upregulated and a gene that encodes serpin (Solyc04g079470) was downregulated. The Arabidopsis homolog of serpin we detected is known to inhibit proteases involved in cell death induction (Lampl et al., 2013; Lema Asqui et al., 2018). Hence, it is possible that during the *Sw-5b*-mediated response, tomato serpin is downregulated allowing the pro-cell death proteases to induce a hypersensitive response in response to TSWV infection.

Protein degradation processes play important roles during defense (Linden and Callis, 2020). At 72 hours post-mechanical inoculation of TSWV, a gene encoding the U-box protein (Solyc04g008100) was upregulated and two genes that encode F-box proteins (Solyc05g055870 and Solyc11g006740) were downregulated (Table 4). Interestingly, Solyc05g025820, a gene that encodes protein kinase was significantly downregulated (Table 4). A homolog of this kinase, Arabidopsis RIPK/PBL14 is known to phosphorylate RIN4 and activate RPM1 NLR-mediated immunity against *P. syringae* expressing avrB and avrRpm1 effectors (Liu et al., 2011). Given the positive regulatory role of this gene in NLR-mediated bacterial immunity, it will be interesting to test how it functions as a negative regulator in *Sw-5b* NLR-mediated resistance against the viral pathogen TSWV. Another interesting gene that is downregulated encodes DCL2b (Solyc11g008540), a dicer-like (DCL) protein. DCLs play an important role in RNA silencing. In tomato, DCL2b is required for the generation of 22-nt small RNAs and defense against *Tomato mosaic virus* (ToMV), *Tobacco mosaic virus* (TMV) and *Potato virus X* (PVX) (Wang et al., 2018a; Wang et al., 2018b). Given DCL2’s function in antiviral defense, it is interesting that DCL2b is downregulated during *Sw-5b*-mediated resistance to TSWV infection.

### Validation of RNA-seq results of some selected genes

To validate the RNA-seq data, we performed RT-qPCR analysis of selected DEGs. We observed a similar trend in expression of selected genes between RT-qPCR and RNA-seq with some difference in magnitude (Fig. 3). Furthermore, the fold change in gene expression observed between RNA-seq and RT-qPCR were strongly correlated (r^2^ = 0.74).

**Figure 3.**
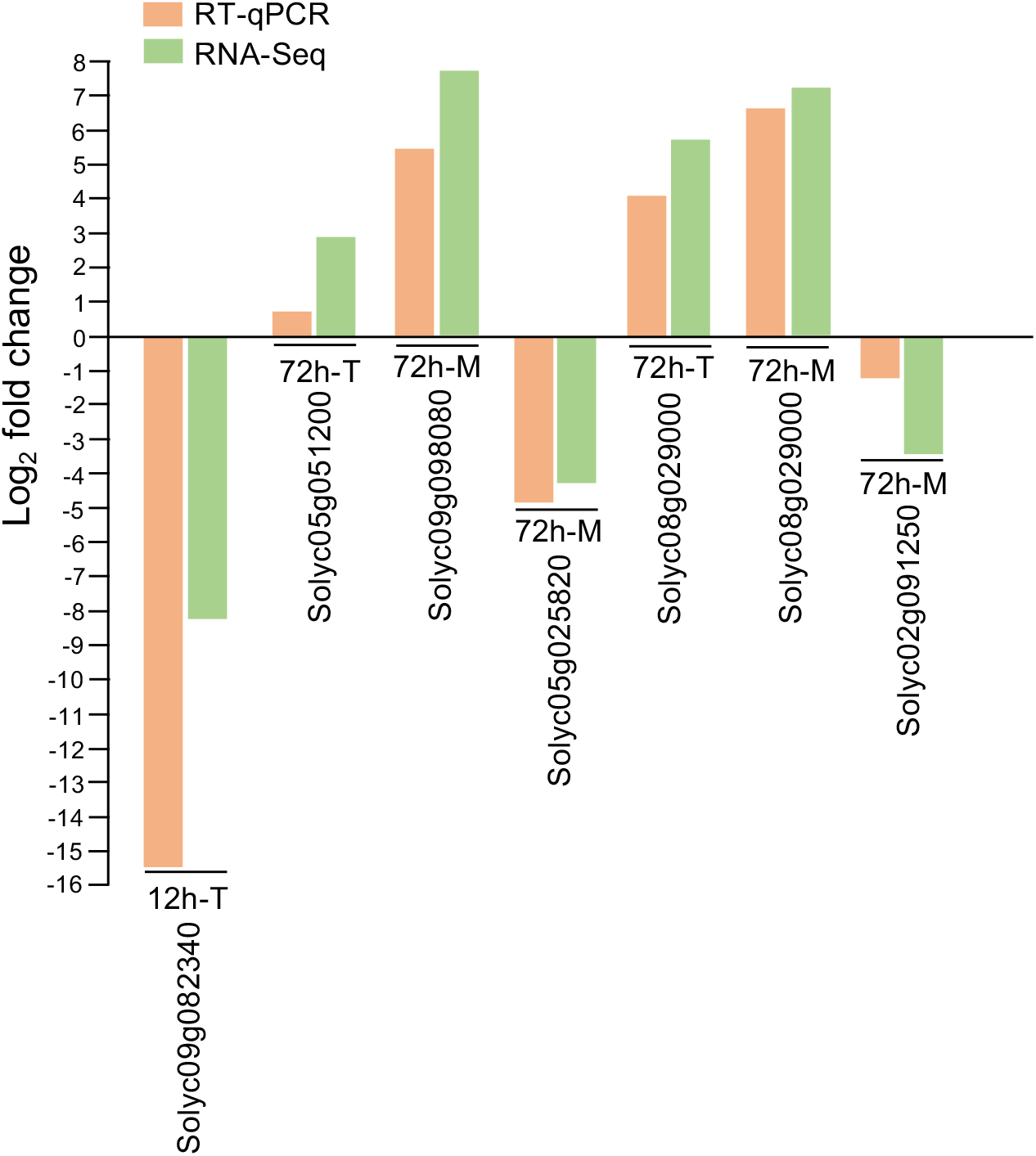
Comparison of RT-qPCR with RNA-seq for selected differentially expressed genes. Expression (log_2_ fold change) of selected genes quantified by RT-qPCR shows the same trend as RNA-seq. See supplementary Table S1 for annotation of genes shown in the graph. T, thrips inoculation; M, mechanical inoculation; h, hours post-inoculation.

### Differences in *Sw-5b* transcriptome changes in response to thrips and mechanical inoculation of TSWV is not due to differences in virus titer

We hypothesized that the earlier response that we observed with thrips inoculations of *Sw-5b* plants may be due to faster or more efficient viral replication in thrips inoculated leaves. Therefore, we determined virus titer by qRT-PCR of the TSWV N gene in plant materials that were used for RNA-seq analysis. We detected TSWV both in thrips and mechanically inoculated leaves from susceptible and resistant lines as early as 12 hours post-inoculation (Fig. 4). Both thrips and mechanical inoculation methods resulted in a significant increase in virus titer in the susceptible line at 72 hours post-inoculation, but virus titers remained low in all time points in the *Sw-5b* resistant line with both inoculation methods (Fig. 4). The abundance of TSWV N gene transcripts were not significantly different in thrips and mechanical inoculated resistant plants. These results suggest that differences in gene expression profiles that we observed with different inoculation methods are not due to differential rates of virus replication.

**Figure 4.**
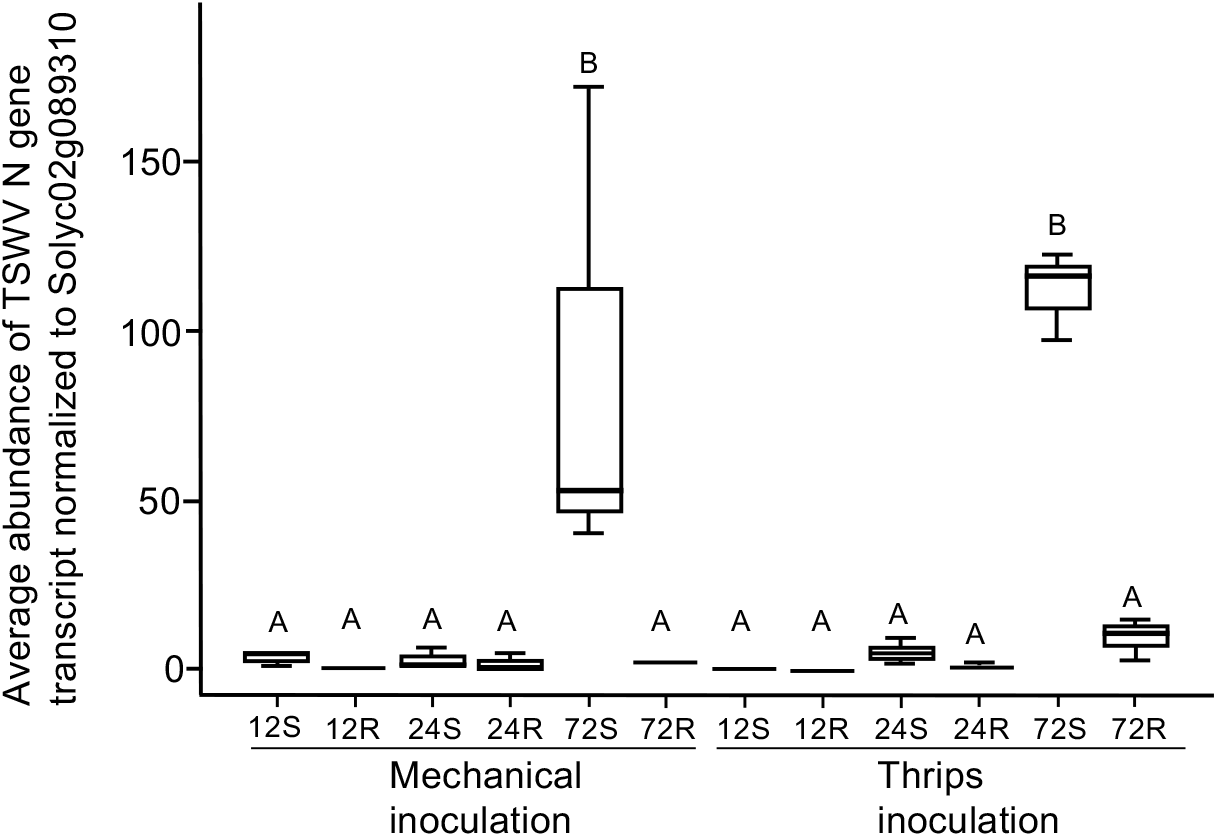
No significant TSWV titer difference in mechanical and thrips inoculated resistant and susceptible tomato plants. Quantification of TSWV by RT-qPCR of resistant (R) and susceptible (S) tomato lines that were infected by mechanical or thrips at 12, 24, 72 hours post inoculation.

## CONCLUSIONS

Our findings revealed that the *Sw-5b* NLR immune receptor induces earlier transcriptome responses to thrips-mediated inoculation of TSWV (12 and 24 HPI) compared to mechanical inoculation (72 HPI). Since activation of the *Sw-5b*-mediated defense requires recognition of TSWV NSm (Chen et al., 2016; Zhu et al., 2017), and this non-structural protein is produced following virus replication (Zhu et al., 2019), we hypothesized that earlier response in the thrips inoculated treatment could be because TSWV replicated more quickly thus NSm is produced earlier than in the mechanical inoculation. However, RT-qPCR analysis of the plant materials used for the RNA-seq library preparation indicated no difference in the virus titer between mechanical and thrips inoculated samples. Thus, some yet to be identified mechanism(s) associated with thrips inoculation of TSWV promotes rapid response of *Sw-5b*.

We anticipated more DEGs at earlier time points compared to the one and fifteen DEGs that we detected at 12 and 24 HPTI, respectively. It is possible that the TSWV replication and synthesis of NSm may require some time before *Sw-5b* recognizes NSm and induces the transcriptome changes. Alternatively, more synchronous induction of infection in many plant cells is required to capture the changes that occur early during infection. While a small number of the DEGs (26) were shared across the inoculation methods, the higher number of DEGs found only in mechanical (69) or thrips (49) inoculation suggesting that the *Sw-5b* NLR uses different mechanisms to provide resistance against TSWV depending on the method of inoculation. In thrips inoculated plants we specifically observed upregulation of genes known to play a role in defense such as a CC-NLR, receptor-like kinases, subtilisin-like protease and transcription factors, ERF1 and CBP60. In mechanically inoculated plants we observed FAD2-9, F-box proteins, homolog of Arabidopsis RIPK/PBL14 kinase, DCL2b and transcription factors, ERF017 and WRKY75.

In conclusion, our findings described here provide new insight into the genes that are differentially regulated in *Sw-5b*-mediated resistance and provide a foundation for future functional analyses to determine the roles of these genes in *Sw-5b*-mediated immune signaling.

## MATERIALS AND METHODS

### Plant and virus material

Near-isogenic TSWV susceptible tomato line ‘Santa Clara’ and TSWV resistant line ‘CNPH-LAM 147’ were described previously (Dianese et al., 2010; Hallwass et al., 2014). Seeds of these lines were planted in individual pots with SunGro professional mix (Agawam, MA) and grown in a growth chamber set at 26°C with 16h day/8h dark, 50% relative humidity, and 300 μEinsteins light intensity. TSWV isolate MR-01 (TSWV ^MR-01^) was collected originally from infected radicchio in Monterey County, CA, flash frozen and stored at −80°C. Complete sequences of this strain can be found in NCBI (https://www.ncbi.nlm.nih.gov/nuccore) using these accession numbers: MG593199 (S RNA); MG593198 (M RNA); and MG593197 (L RNA) (Adegbola et al., unpublished data).

### Thrips colony and maintenance

*Frankliniella occidentalis* from a lab colony originally collected from the Kamiloiki valley on the Hawaiian island of Oahu was reared on green pods of *Phaseolus vulgaris* as previously described (Ullman et al., 1992; Bautista, 1993). Thrips inoculations were standardized and optimized by collecting thrips larvae less than six hours post-eclosion, then placing them on either TSWV^MR-01^ infected *D. stramonium* for a 24-hour acquisition access period (AAP). After the AAP, thrips were reared on green bean pods and at 24-hr post adult eclosion were used for thrips inoculations. Noninfected control thrips were created using the same strategy, except they were given acquisition access to non-infected *Datura stramonium*.

### Mechanical inoculation of TSWV

For mechanical inoculation of tomato lines, TSWV^MR-01^ inoculum was prepared by grinding one gram of *Datura stramonium* TSWV infected leaf tissue in 15 ml of buffer (0.1M potassium phosphate, 1 mM sulphite, 1% celite, pH 7) with a mortar and pestle on ice. The equal amount of resulting sap was rub inoculated onto leaves using a pestle. Ten minutes after the inoculations, the inoculated leaves were rinsed with distilled water and maintained in the growth chamber with conditions as described above until the samples were collected for RNA extraction. The presence of TSWV^MR-01^ was confirmed by enzyme-linked-immunosorbent assay (ELISA) following the manufacturer’s protocol (Agdia, IN).

### TSWV inoculation through thrips

Single tomato plants and thrips were placed in cages made up of two-32-ounce plastic cups (Lancaster, PA) placed on the pot with the bottom cut out and covered with no-thrips insect screens (BioQuip, CA) (Supplementary Fig. S1). The pot was covered using parafilm to prevent thrips movement into the soil. Ten thrips (5 females and 5 males) were placed on leaf 1 and leaf 2 on each plant, and the cage closed. Thrips remained on the plants until samples were collected for RNA extraction.

### Tissue collection for RNA sequencing

Samples were collected at 0, 12, 24 and 72 hours post mechanical and thrips inoculation. At each collection time, the two inoculated leaves from each plant (n=10) were collected from each treatment: viruliferous thrips (VT), non-viruliferous thrips (NVT) (mock), mechanical inoculation with sap from virus-infected plants (VS), mechanical inoculation with sap from non-virus infected plants (NVS) (Figure 1D). TSWV infection of the positive controls that were inoculated with VT and VS were used to verify inoculation success using ELISA. The no thrips control was used to test for cross-contamination during the biological experiments. The collected tissue samples were flash-frozen in liquid nitrogen and stored at −80° C. Additional confirmation of TSWV presence, and concentration was assayed using RT-qPCR with primers that anneal to part of the *N* gene of TSWV (Supplementary Table S2). RT-qPCR was performed for three biological replicates and six technical replicates/biological replicates using SYBR green mix on Bio-Rad CFX96 machine with the following conditions: 95 °C for 30 sec, followed by 39 cycles of 95 °C for 10 sec, 55 °C for 10 sec, and 60 °C for 20 sec.

### RNA extraction and RNA-seq library preparation

Tissue samples collected as described above were used for the preparation of RNA-seq libraries using the protocol described in (Nagalakshmi et al., 2010). Briefly, mRNA was isolated using oligo (dT) magnetic beads (Invitrogen, CA), treated with DNase, first and second-strand cDNA synthesis, followed by fragmentation and addition of barcoded adaptors. A total of 96 barcoded libraries were prepared, pooled, and sequenced using the Illumina HiSeq4000.

### Differential gene expression analysis

The sequencing adaptors and low-quality bases were trimmed from the raw reads using Trimmomatic version 0.36 (Bolger et al., 2014). The high-quality reads were mapped to the tomato cDNA sequences (version 4.1; Sol Genomics, https://solgenomics.net/) using Salmon version 0.8.1 (Patro et al., 2017). Differential gene expression analysis was performed using the generalized linear model (glm) functionality of the edgeR package (Robinson et al., 2010). Tomato genes with at least two fold expression difference between the susceptible and resistant genotypes and False Discovery Rate (FDR) < 0.05 were considered differentially expressed.

### Quantitative RT-PCR

Gene-specific primers were designed using Primer3Plus for amplification of 93-180 bp fragments from each target gene (Supplementary Table S2). The F-box gene was used to normalize the data (Liu et al., 2012). Total RNA was extracted from frozen leaf material using TRIzol® (Invitrogen, CA) according to manufacturer’s instructions. RNA was treated with DNase I (Invitrogen, CA) followed by cDNA synthesis using Verso cDNA Synthesis kit (Thermo Fisher Scientific, MA). Three biological replicates were run on Bio-Rad CFX96 machine using the following conditions: 95 °C for 30 sec, followed by 39 cycles of 95 °C for 10 sec, 55 °C for 10 sec and 60 °C for 20 sec. The gene expression fold change was calculated using the 2(−ΔΔCT) method (Livak and Schmittgen, 2001).

## ACKNOWLEDGEMENTS

This work was supported by National Science Foundation grant IOS-1339185 (to SPD-K and DEU). We thank Dr. Renato Resende, University of Brasilia, who contributed to development of the tomato isogenic lines used in our investigation, Dr. Sulley Ben-Mahmoud who provided technical assistance for RT-qPCR aspects of our work, and to Dr. Robert Gilbertson who graciously read and edited early drafts of the manuscript.

---

**Supplementary Figure S1.**
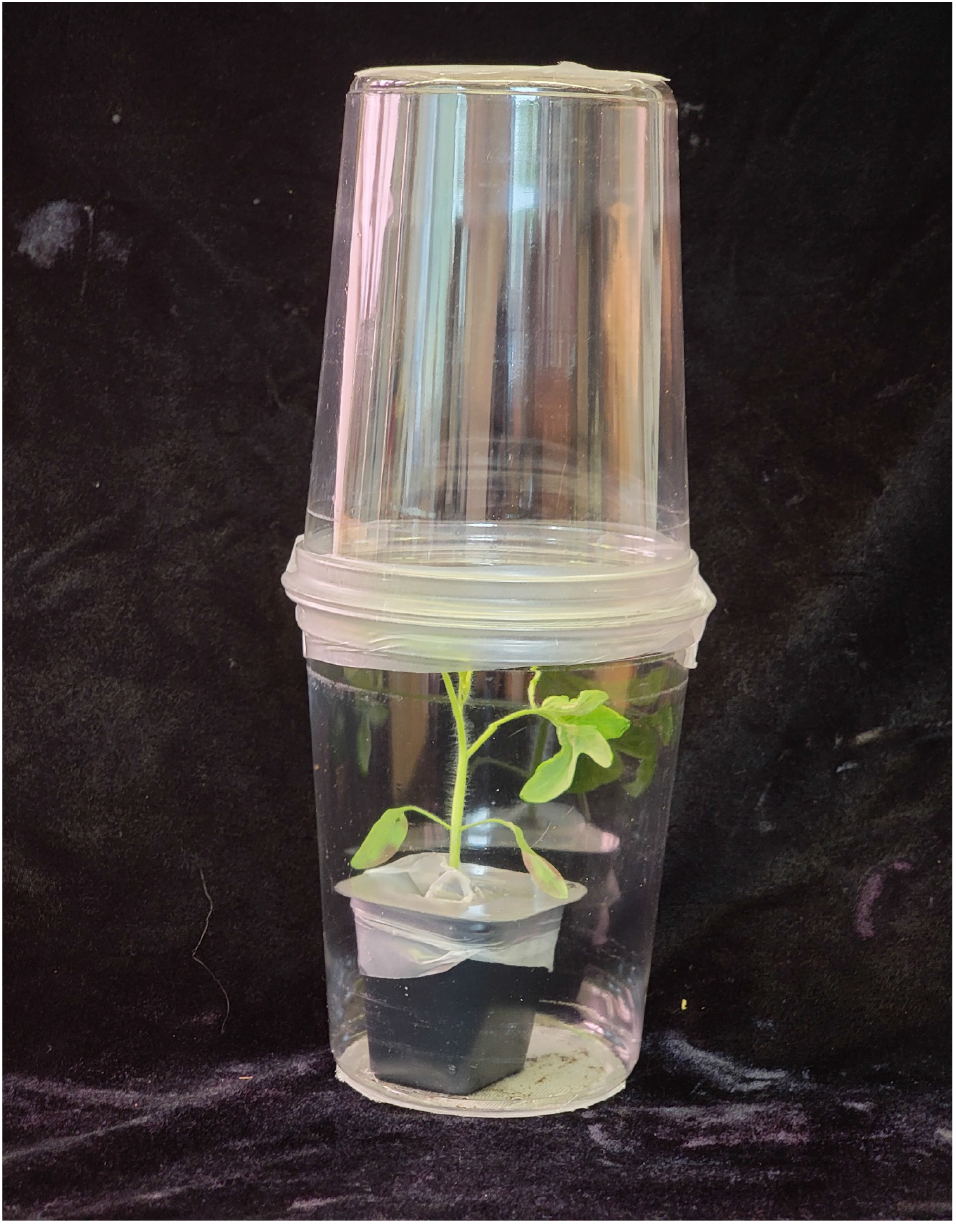
Cage used for inoculation of TSWV through thrips.

