## Supplementary Table S2 for "The *Sw-5b* NLR immune receptor induces earlier transcriptional changes in response to thrips-mediated inoculation of *Tomato spotted wilt orthotospovirus* compared to mechanical inoculation"

**Supplementary Table S2.** Primers used in this study.

| Primer | Primer sequence (5'-3') | Purpose |
|---|---|---|
| Solyc02g089310 (F-Box) | F: GGAACTCACAAACGCCTATTT | RT-qPCR |
| | R: ATCTTGGAGGCATCCTGCTTAT | |
| Solyc09g082340 (Vicilin) | F: ACCAGAAAATCCGCGGTAAC | RT-qPCR |
| | R: AGATTCGAGTTTCCGGTTGC | |
| Solyc09g098080 (Glycosyltransferase) | F: ACCAACAAGTTACAGCGACTAT | RT-qPCR |
| | R: TGTGGTGCCCATCCTATTATTT | |
| Solyc05g025820.3.1 (Protein kinase family protein) | F: CACAAACGAAGTCCTCAAAGGG | RT-qPCR |
| | R: TGGAAATCATTGTGCAATGGTG | |
| Solyc08g029000 (Lipoxygenase) | F: ACGGCAGCGTTCATTTTGTC | RT-qPCR |
| | R: ATTGCGCAATGGTTCTGGTG | |
| Solyc02g091250 (Mediator of RNA polymerase II transcription subunit ) | F: GACCACCTTCAAATTGGATGGG | RT-qPCR |
| | R: AACATCCCCATAGCCAAAGC | |
| Solyc05g051200 (ERF1) | F: GGCCATGGGGTAAATATGCATC | RT-qPCR |
| | R: AGACCAAGGACCCCTCATTG | |
| TSWV – N ORF | F: GCTTCCCACCCTTTGATTC | RT-qPCR |
| | R: ATAGCCAAGACAACACTGC | |
