## Supplementary figures and images for "The *Sw-5b* NLR immune receptor induces earlier transcriptional changes in response to thrips-mediated inoculation of *Tomato spotted wilt orthotospovirus* compared to mechanical inoculation"

### Supplementary Figure S1

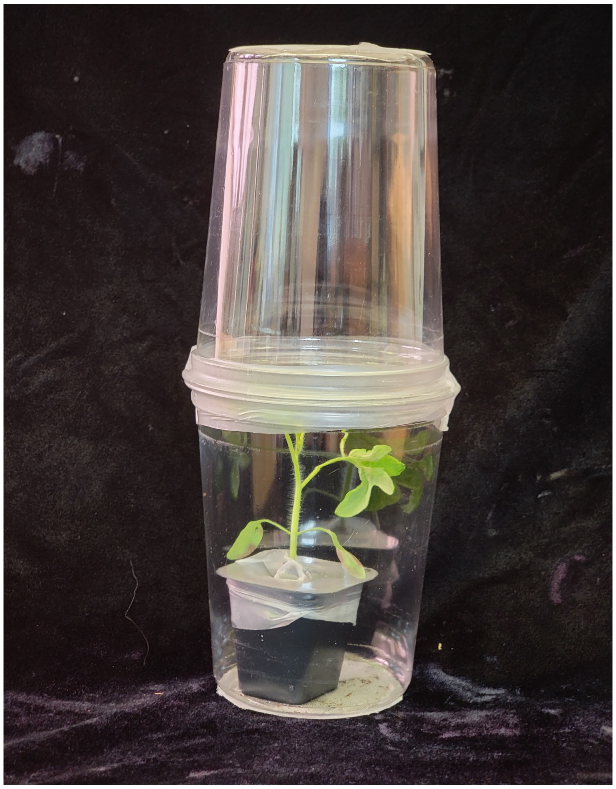

**Supplementary Figure S1.** Cage used for inoculation of TSWV through thrips.
